# In-Chamber Sublimation: A Practical Approach for Mitigating Ice and Curtaining in Cryo-Electron Tomography Lamellae Preparation

**DOI:** 10.64898/2026.03.12.711158

**Authors:** Amy L. Bondy, Genís Valentín Gesé, Thomas Thersleff, B. Martin Hällberg

## Abstract

Surface ice contamination is a persistent challenge in cryo-electron tomography (cryo-ET) workflows, where it can obscure regions of interest and contribute to curtaining artefacts during focused ion beam (FIB) milling. We demonstrate using high-pressure frozen yeast cells that a sublimation step within the scanning electron microscope (SEM) chamber before lamella milling visually removes surface ice and reduces sample roughness without detectable devitrification. While sublimation has been widely applied in cryo-SEM and volume imaging, it is not common on cryo-ET samples due to concerns about devitrification. Using tomographic reconstructions, we show that controlled sublimation improves lamella quality by reducing surface roughness and minimizing curtaining without compromising sample vitrification. Furthermore, subtomogram averaging of the 80S ribosome confirmed lamellae quality are preserved after sublimation. This approach offers a practical refinement to existing cryo-ET preparation protocols, requiring no additional instrumentation or workflow modifications.

## 1. Introduction

Cryogenic electron microscopy (cryo-EM) techniques have become essential tools in structural and cellular biology, enabling high-resolution imaging of biological specimens in their near-native, vitrified state. Among these methods, cryo-focused ion beam scanning electron microscopy (cryo-FIB-SEM) is widely used for site-specific milling of thin lamellae, which are then imaged using cryo-electron tomography (cryo-ET) [1,2,3]. Cryo-ET, in particular, offers three-dimensional imaging of cells at sub-nanometer resolution, providing critical insights into macromolecular organization in situ. Other cryo volume imaging approaches, such as cryo-serial block-face imaging, rely on similar workflows to access and image the sample’s interior [4]. However, across all these techniques, surface ice contamination remains a persistent challenge, often obscuring features of interest or interfering with sample preparation steps such as ion milling [5,6].

Several strategies have been developed to mitigate surface ice accumulation. These include expensive facility upgrades (e.g., installing low-humidity cleanrooms) [7], investment in specialized instrumentation (e.g., glove boxes, vacuum transfer systems, or in-chamber fluorescence microscopes to reduce handling steps) [5,8], or routine interventions such as filtering liquid nitrogen with cloth or paper filters. In some workflows, large pieces of surface ice are manually removed with a cryo-compatible paintbrush [9]. This method is particularly common for high-pressure frozen (HPF) samples, which often accumulate ice during the mechanical fracturing of sample carriers prior to SEM or FIB imaging. While these measures can help reduce contamination, they each carry limitations. Manual cleaning risks damaging the sample; infrastructure improvements may be cost-prohibitive and not an option for smaller microscopy facilities; and even with careful handling, transfer steps remain a significant source of potential contamination.

Surface ice contamination is particularly problematic for cryo-ET and cryo volume imaging workflows. In cryo-FIB-SEM, a rough or ice-covered surface can lead to curtaining artefacts during milling, compromising lamellae quality [10]. In imaging applications, ice obscures cellular features, making regions of interest inaccessible or uninterpretable. These challenges are especially acute in workflows involving correlative fluorescence microscopy, where multiple transfers and extended handling can increase the risk of contamination [5].

Sublimation, or controlled thermal desorption of amorphous ice under vacuum, has long been used in cryo-SEM to remove superficial ice and reveal underlying ultrastructure [10,11,12,13]. In some FIB-milling and SEM imaging applications, sublimation has also been used to reduce surface roughness and improve milling performance [10,14].

Importantly, sublimation can be performed either in an external chamber or directly within the cryo-SEM, requiring no additional hardware. However, despite its utility, sublimation has been avoided in workflows that include cryo-TEM imaging, due to concerns that warming the sample—even under vacuum—could lead to devitrification of vitreous ice [10,14].

In this study, we examine these concerns by systematically evaluating sublimation under controlled conditions. We demonstrate using high pressure frozen yeast cells as a test sample that sublimation, when carefully performed within the SEM chamber under controlled conditions, does not induce detectable devitrification and can be safely integrated into cryo-ET workflows. Moreover, sublimation is highly effective at removing surface ice contamination and reducing curtaining artefacts, leading to improved lamellae and tomogram quality, capable of achieving high resolution during subtomogram averaging. Our findings suggest that sublimation, long used in cryo-SEM for surface cleaning, can now be re-evaluated as a viable and accessible tool for cryo-ET sample preparation.

## 2. Methods

All experiments were performed on high-pressure frozen yeast “waffle” samples in humidity-controlled rooms. All samples for both the sublimation experiments and the control experiment were handled and stored in the same manner throughout the entirety of the experiments. The rooms in the cryo-EM facility in which the following steps were carried out are humidity-controlled to 10% humidity: high pressure freezing, auto grid clipping, cryo-confocal loading and imaging, TEM station loading, and sample storage. The room where the Aquilos 2 cryo-FIB-SEM loading/unloading was carried out is humidity-controlled to 30% humidity. Fig. 1 shows the general workflow used in this paper, denoting when sublimation was carried out in the workflow.

**Figure 1.**
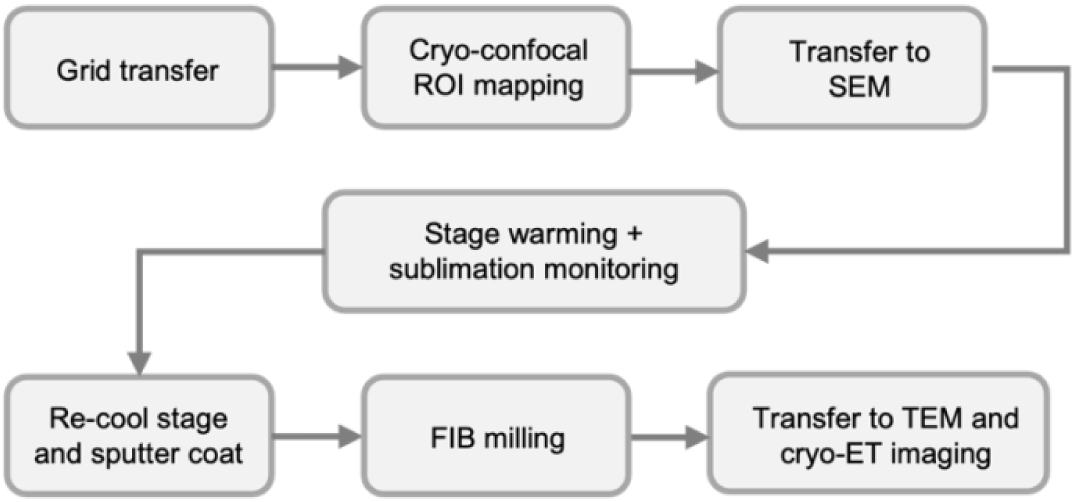
A schematic diagram of the general workflow used in this paper, highlighting the several transfer steps needed for the fluorescence-targeted cryo-ET workflow, and at which stage in the workflow sublimation was carried out.

2.1. *Yeast cells sample preparation*

Yeast cells (BY4741 RPM2-GFP his3MX6 strain from the Yeast GFP Collection [15]) tagged with green fluorescent protein (GFP) were grown on an agar plate at room temperature. Following sufficient colony growth (approximately 24 hours), the plate was transferred and stored at 4°C until used. Cells were harvested from the plate by transferring a centimeter scoop (amount is approximate) into a 0.15 mg/mL solution of Evans Blue in phosphate buffer to create a highly viscous cell suspension solution. The cells were incubated for 5 minutes just prior to grid preparation.

2.2. *High-pressure freezing*

Grids were prepared using the ‘waffle method’ [16]. In brief, the flat side of 3 mm planchets were polished with sandpaper and then coated with 2 µL of silicone solution in isopropanol (Serva, product no. 35130), which was allowed to evaporate fully before use. TEM grids (400×100 mesh, Cu, Agar Scientific) were glow discharged on both sides at 35 mA for 2 minutes. The waffle grid was assembled in the high-pressure freezer (HPF) as follows: planchette hat (flat side up), glow-discharged Cu grid, 2 µL of cell suspension, planchette hat (flat side down). Excess sample was wiped off with Whatman filter paper (type 1). This assembly was inserted into a Leica EM ICE (Leica, Wetzlar, Germany) and high-pressure frozen under conditions of 325 milliseconds and pressure of 2200 bar. The waffle was then disassembled under liquid nitrogen and clipped into an autogrid.

2.3. *Correlative cryo-confocal imaging*

Frozen autogrids were imaged by cryo-confocal microscopy using the sp8 microscope (Leica) equipped with a cryo-stage and 100x/0.9 NA objective lens. Initially the grids were imaged in brightfield mode using transmitted light to identify regions of interest with densely packed cells. An overview tile set of the entire grid was collected, and the tiles were merged for later exportation. In confocal mode, the 488 nm and 640 nm channels were used to image GFP (499nm – 573nm, Hyb2 detector, 1.5% laser power) and Evans Blue (648nm – 750nm, Hyb3 detector, 0.1% laser power), respectively. Z-stacks were collected from two regions of interest (z step = 0.4 µm, 400×400 scan speed, resolution = 1024×1024, zoom = 3.00, line averaging = 2). After confocal imaging the maximum projections of each z-stack were processed to assist with the correlation. Images (transmitted light merged tile set, z-stacks, and z-stack maximum projections) were imported into Maps version 3.31 (Thermo Fisher Scientific) for correlation.

2.4. *Cryo-FIB lamellae preparation*

Note: Three sets of experiments were conducted on separate days, so that different in-chamber sublimation conditions could be followed. All procedures followed essentially the same protocol however for imaging and lamellae preparation, which is described below.

Following cryo-confocal imaging, waffle grids were placed into a 45° pre-tilted cryo-FIB-SEM shuttle and inserted into the pre-cooled Aquilos 2 cryo-FIB-SEM (Thermo Fisher Scientific) for imaging and lamellae preparation. Initially, overview tile sets were collected in Maps. For the control sample without sublimation, the sample was then sputter coated with Pt (30 mA, 15 s). For the two experiments that included a sublimation step, sublimation was carried out prior to sputter coating. For in-chamber sublimation, the flow rate of cooling N_2_ gas was lowered (∼145 mg/s) to increase the cryo-stage temperature. The N_2_ gas flow was lowered step-wise (starting at 200 mg/s, then lowered to 170 mg/s, then ∼ 145 mg/s), decreasing flow once the temperature started to stabilize at a specific flow rate. This step-wise decrease in N_2_ flow, and subsequently stage warming, was important to ensure that 1) the cryo-shield temperature always remained cooler than the cryo-stage to trap sublimated water, 2) the cryo-stage did not warm too quickly and severely exceed the minimum sublimation temperature leading to freeze-etching [17], and 3) once the sublimation temperature was reached, the temperature could be approximately maintained.

2.4.1. *Slow-sublimation experiment*

For the slow-sublimation experiment, the stage temperature was quickly warmed to −118°C using the N_2_ gas flow rates listed above (always keeping the cryo-shield temperature cooler than the cryo-stage temperature) and then slowly warmed to −104°C over a period of two hours. The extent of surface ice was observed with SEM images collected every 5 minutes and ion images collected every 10-15 minutes. Based on these images, sublimation was observed to begin after approximately 70 minutes (−113°C) which is denoted as T_s_ = 0 in the text. Sublimation was terminated at T_s_ = 55 minutes because almost all the surface ice was gone at this timepoint. The stage temperature was then quickly cooled to −175°C by increasing the N_2_ gas flow rate (200 mg/s), and the grid was sputter coated with Pt (30 mA, 15 s).

2.4.2. *Fast-sublimation experiment*

For the fast-sublimation experiment, the stage temperature was quickly warmed to −120°C (always keeping the cryo-shield temperature cooler than the cryo-stage temperature) and then warmed to −97°C over a period of 35 minutes. Sublimation was terminated after 35 minutes, whereupon the stage temperature was quickly cooled to −175°C by increasing the N_2_ gas flow rate (200 mg/s), and the grid was sputter coated with Pt (30 mA, 15 s).

2.4.3. *Surface ice monitoring*

Surface ice was dynamically monitored during sublimation by acquiring SEM images at regular intervals. Ice content was assessed by manual visual inspection during the experiment, and this assessment was used to define the sublimation endpoint. Relative ice content was subsequently quantified post-experiment, on the assumption that surface ice directly correlates with the root-mean-square (RMS) roughness of preprocessed images (Table S1). This metric is analogous to the *S_q_* surface roughness parameter [18], applied here to the image intensity domain as a proxy for surface texture, with *S_q_* decreasing as surface smoothing reduces intensity edges and local contrast. Reproducible, comparable *S_q_* values across timepoints were ensured by applying an identical preprocessing pipeline to every image: images were first normalized to [0, 1], the background was estimated by morphological opening with a disk-shaped structuring element (radius 50 µm) and subtracted, and the result was denoised with a Gaussian filter (σ = 630 nm) before computing *S_q_*. Standard errors were estimated via spatial block bootstrapping, dividing each preprocessed image into non-overlapping 256 x 256 pixel tiles, computing *S_q_* per tile, and reporting the standard error of the mean across tiles.

2.4.4. *Milling procedure for all experiments*

Following sputter coating, a new tile set was captured in Maps and used for correlation with the confocal data. The grid was then put in the deposition position and coated with platinum GIS for 2 minutes. The waffle method milling procedure for *E. coli* was then followed (see Kelley et. al. [16] for stage positions, angles, and milling currents used for each step). Briefly, two trenches were milled for each lamella site at the orthogonal ion beam position. Clean-up of the bottom trenches was conducted at higher milling angles (∼26°) and then the milling angle (20° or 16°, depending on the lamella site). Using AutoTEM Cryo 2.4 the preparation step, which includes eucentric tilting, setting the milling angle, collecting a reference image, and lamella placement, was done for each lamella site. Each lamella was then milled using the Rough Milling, Medium Milling, Fine Milling, and Finer Milling parameters cited above. Lamellae were manually polished using cleaning cross-section patterns and a milling current of 30 pA. Following final lamellae thinning, grids were sputter coated with Pt (7 mA, 10 s, 0.10 mbar) to reduce charging before unloading from the cryo-FIB-SEM.

2.5. *Cryo-ET collection*

Tilt-series for lamellae from the slow-sublimation experiment were collected with Tomography 5.22 using a Titan Krios (Thermo Fisher Scientific) equipped with a Gatan K3 BioQuantum energy filter (counting mode, energy filter slit width = 10 eV). Tilt series were collected in superresolution mode yielding a pixel size of 1.34 Å, with a nominal magnification of 33,000, target defocus of –6 µm, tilt range of −40° to +40° from the milling angle, and 100 e/Å^2^ total dose per tilt-series. Collection was performed using the dose symmetric collection scheme with 2° tilt increments. A total of 16 tilt series were collected.

2.6. *Cryo-ET processing*

Tilt images were binned 2x (2.68 Å/px) and frame aligned using Warp [19] with a 1×1×6 model. Tilt images with an estimated resolution worse than 15 Å were discarded. Tilt series were binned 4x (10.72 Å/px), aligned via patch tracking (3×2 patches) and used for tomogram reconstruction (SIRT) in AreTomo2 [20]. Lamella thickness measured from the reconstructed tomograms was 287 ± 33 nm (mean ± SD).

2.7. *Subtomogram averaging*

Tomograms for template matching were reconstructed in WARP 1.0.9 [19], using alignments produced by AreTomo2, at a pixel size of 16.08 Å. Template matching was performed using pyTME 0.3.4a [21], with EMD-3228 used as the initial reference.

Angular sampling was set to 8°, and the reference was low-pass filtered to 60 Å. WARP XML metadata were provided to pyTME to enable dose weighting and CTF correction during matching, using a spherical aberration of 2.7 mm, an acceleration voltage of 300 kV, and an amplitude contrast of 0.07. Candidate particles were selected using a minimum inter-particle distance of 160 Å and a minimum cross-correlation score of 0.20. To restrict particle selection to the cytosolic compartment, binary masks were manually generated in UCSF ChimeraX [22] using the ArtiaX plugin [23] and supplied to pyTME.

Subtomograms were extracted in WARP at a pixel size of 10.72 Å with a box size of 72 pixels. Initial dataset cleaning was performed by 3D classification in RELION 3.1 [24], and particles from the highest-resolution classes were selected for further processing. These particles were subjected to iterative subtomogram refinement in M, using a three-step refinement protocol (Table S2). Based on the refined particle alignments obtained in M, improved tomograms were reconstructed in WARP and used for a second round of template matching. This round employed identical matching parameters as described above, except that the refined subtomogram average from M was used as the reference. Particles from the second template-matching round were extracted in WARP at a pixel size of 5.36 Å with a box size of 144 pixels. The extracted particles were refined in RELION 3.1 using 3D auto-refine, followed by 3D classification without alignment to remove residual bad particles. Ultimately 2,738 particles from 10 tilt series were used. Particles from the best-resolved classes were subsequently subjected to a final round of subtomogram refinement in M using the same refinement strategy. Finally, particles were re-exported from M at a pixel size of 2.68 (bin 1) with a box size of 288 pixels for a final round of gold-standard 3D auto-refine in RELION 3.1.

To quantify the effect of the sublimation treatment on lamella quality, a Rosenthal-Henderson plot [25] was produced (Fig. 5). After the final RELION 3.1 refinement, random subsets of particles with varying sizes were selected ensuring equal representation from each of the two independently refined half-datasets. For each subset, two half-maps were reconstructed, and their Fourier shell correlation (FSC) was calculated using RELION 3.1. The Rosenthal-Henderson B-factor was fitted using the equation: 1/𝑑2 = (2/𝐵)𝑙𝑛(𝑛𝑢𝑚𝑏𝑒𝑟 𝑜𝑓 𝑝𝑎𝑟𝑡𝑖𝑐𝑙𝑒𝑠) as implemented in Zivanov et al. [26]. Local resolution estimates were computed using relion_postprocess as implemented in RELION 3.1.

## 3. Results and Discussion

3.1. *Sublimation decreased lamellae roughness*

To evaluate how sample sublimation benefits FIB-milling, high-pressure frozen yeast samples with visible surface ice were selected for lamellae preparation. Sublimation was initiated by gradually increasing the cryo-stage temperature under SEM vacuum while monitoring the process in real time. Visual inspection via SEM and ion beam imaging at regular intervals allowed termination once surface ice was no longer detectable (Fig. 2a–l). Roughness calculations performed afterward on the normalized and filtered SEM images (Table S1) quantify the decreasing surface roughness as the surface ice is sublimated away. This real-time monitoring provides an operational advantage over external sublimation setups that lack direct visual control [10].

**Figure 2.**
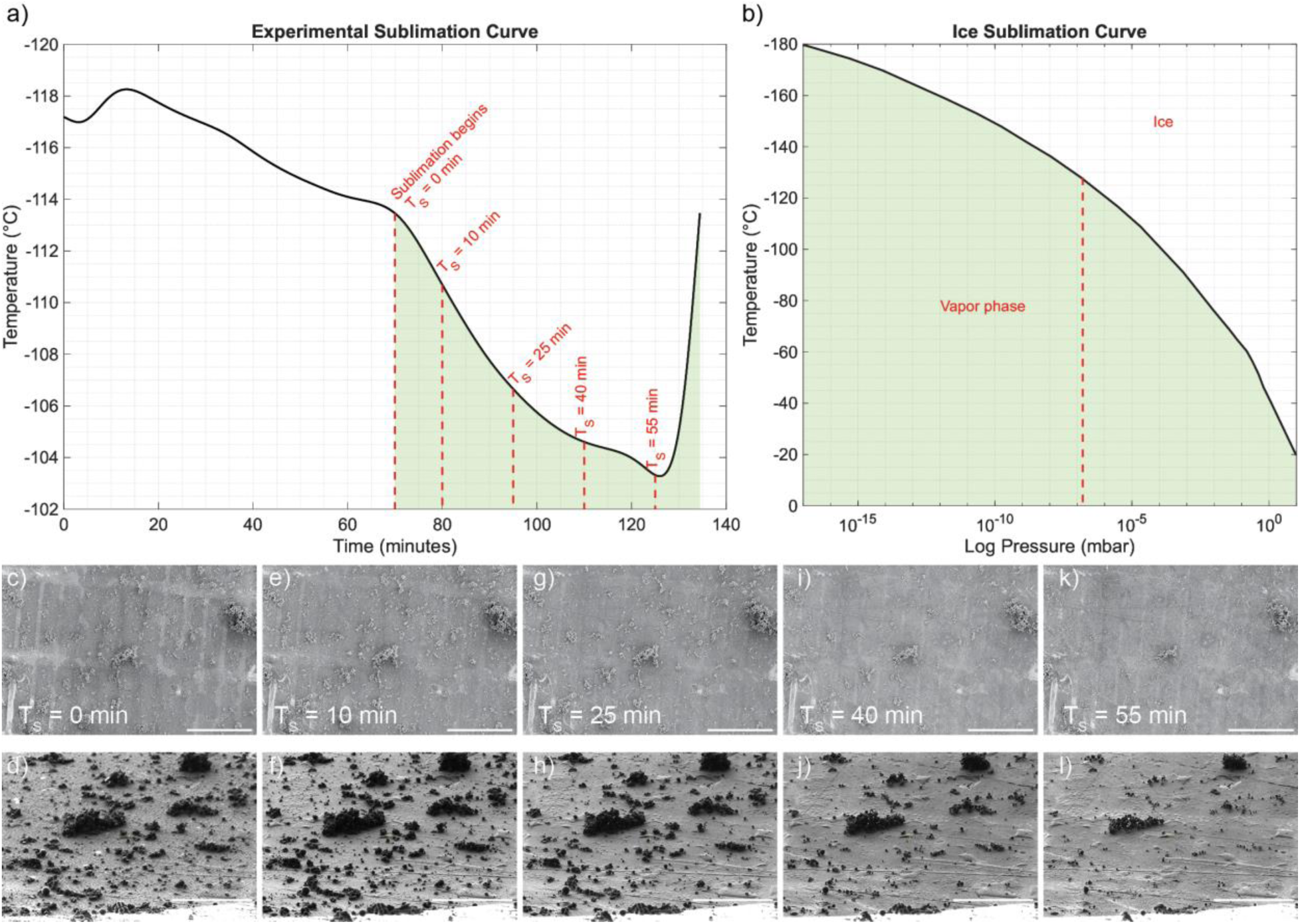
Temperature curves showing sublimation of a) surface ice on the sample (experimental) vs. b) theoretical sublimation curve of ice at low pressure (based on Boës et al. [37]), where the red dashed line shows the actual pressure of the SEM during the experiment. Images c) through l) show the progression of sublimation over time for this experiment at T_s_ = 0 in the c) SEM image and d) ion image; T_s_ = 10 minutes in the e) SEM image and f) ion image; T_s_ = 25 minutes in the g) SEM image and h) ion image; T_s_ = 40 minutes in the i) SEM image and j) ion image, and at T_s_ = 55 minutes in the k) SEM image and l) ion image when sublimation was ceased. The scale bars are 200 µm for the SEM images and 100 µm for the ion images, respectively.

When used prior to FIB milling, sublimation markedly improved lamellae quality. Lamellae from sublimated samples showed smoother surfaces and minimal curtaining artefacts at all stages of milling, including trench milling, rough milling, and thinning (Fig. 3a–c), consistent with the progressive smoothing observed during sublimation (Table S1). In contrast, untreated samples retained a rough surface layer and displayed severe curtaining during milling (Fig. 3d–f). These observations indicate that sublimation reduces surface roughness and facilitates uniform lamellae preparation.

**Figure 3.**
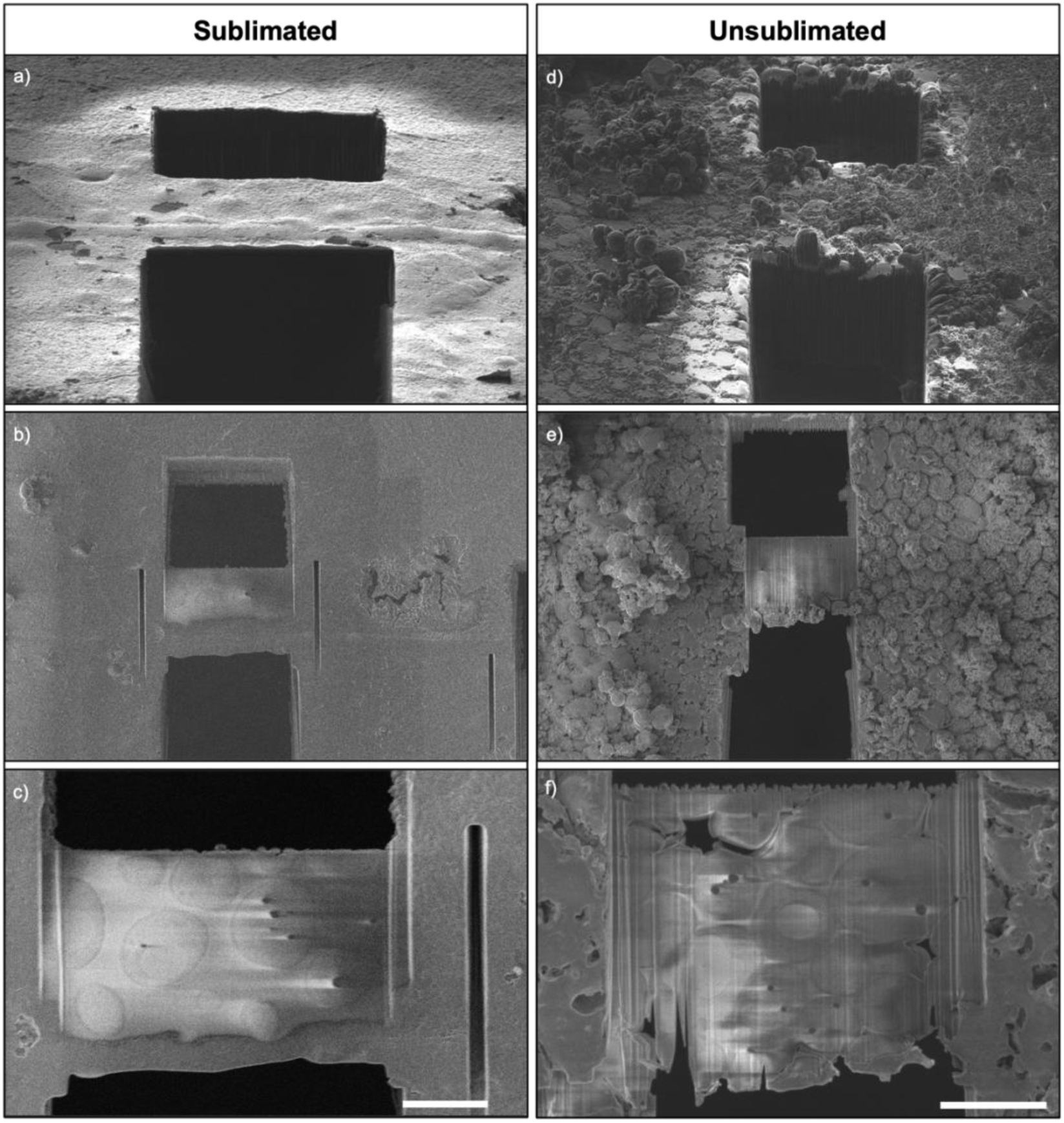
SEM images of HPF frozen yeast lamellae during the milling process. Images a) through c) show a sample that has been sublimated prior to milling, while d) through f) are from an unsublimated sample with extensive surface ice contamination. A) and d) show the lamellae after trench milling (waffle method); b) and e) show after rough milling; and c) and f) show after thinning. Curtaining in the final lamellae is much more severe for the unsublimated sample with a rough surface. The scale bars for final micrographs are 5 µm.

3.2. *Sublimated sample maintains vitreous state*

We next sought to rule out partial devitrification caused by the sublimation procedure, which would render the sample unsuitable for cryo-ET. To assess this, we acquired tilt series from lamellae milled on sublimated samples. All tomograms showed preserved ultrastructure and lacked evidence of crystalline ice [27,28,29,30]. For our primary evidence of the vitreous state of the lamellae, the raw images as well as the aligned tilt series showed no diffraction effects or ordered, crystalline regions (Fig. 4a, c-f) [31,32,33]. This projection-level evidence is the primary indicator of vitrification and tomogram-slice FFTs are provided as qualitative complements. Movies S1–S12 further confirm vitrification across multiple datasets. Further supporting this claim, Fourier transforms of tomogram slices exhibited only diffuse intensity, with no reflections corresponding to cubic (3.1 nm⁻¹) or hexagonal (2.5 nm⁻¹) ice (Fig. 4b).

**Figure 4.**
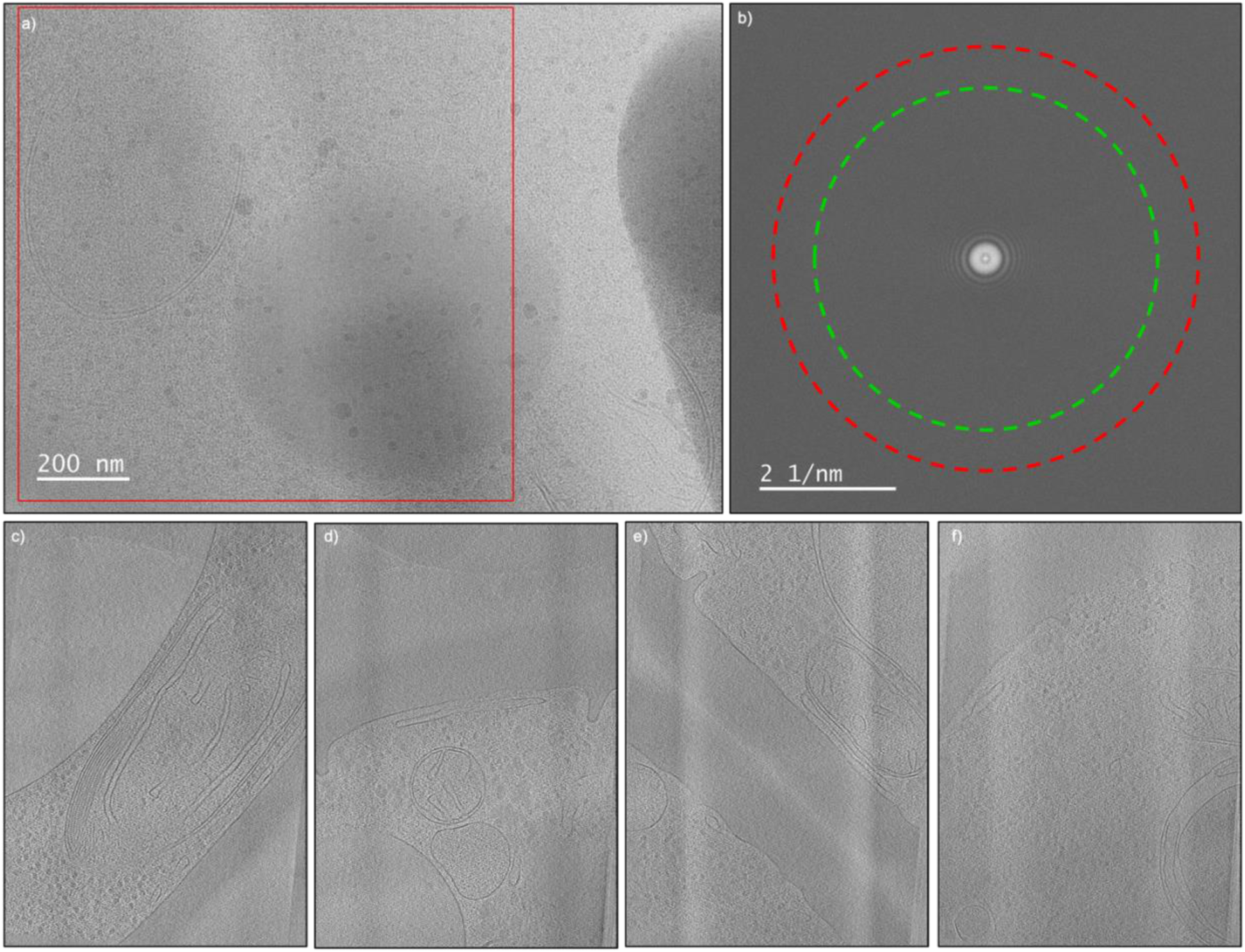
Tomograms of HPF yeast lamellae where the sample was sublimated before milling to remove surface ice show no evidence of crystalline ice. a) Shows an unprocessed tilt image (tilt 20°, pixel size 1.34 Å) and b) the corresponding FFT averaged over the square region marked in red. The FFT shows no evidence of cubic ice (red circle; 3.1 nm^-1^) or hexagonal ice (green circle; 2.5 nm^-1^) which would show as bright spots rather than diffuse rings. Images c) through f) show sections from different tomograms in the same experiment, none of which show evidence of crystalline ice. Movies of all tomograms are in the Supporting Information.

**Figure 5.**
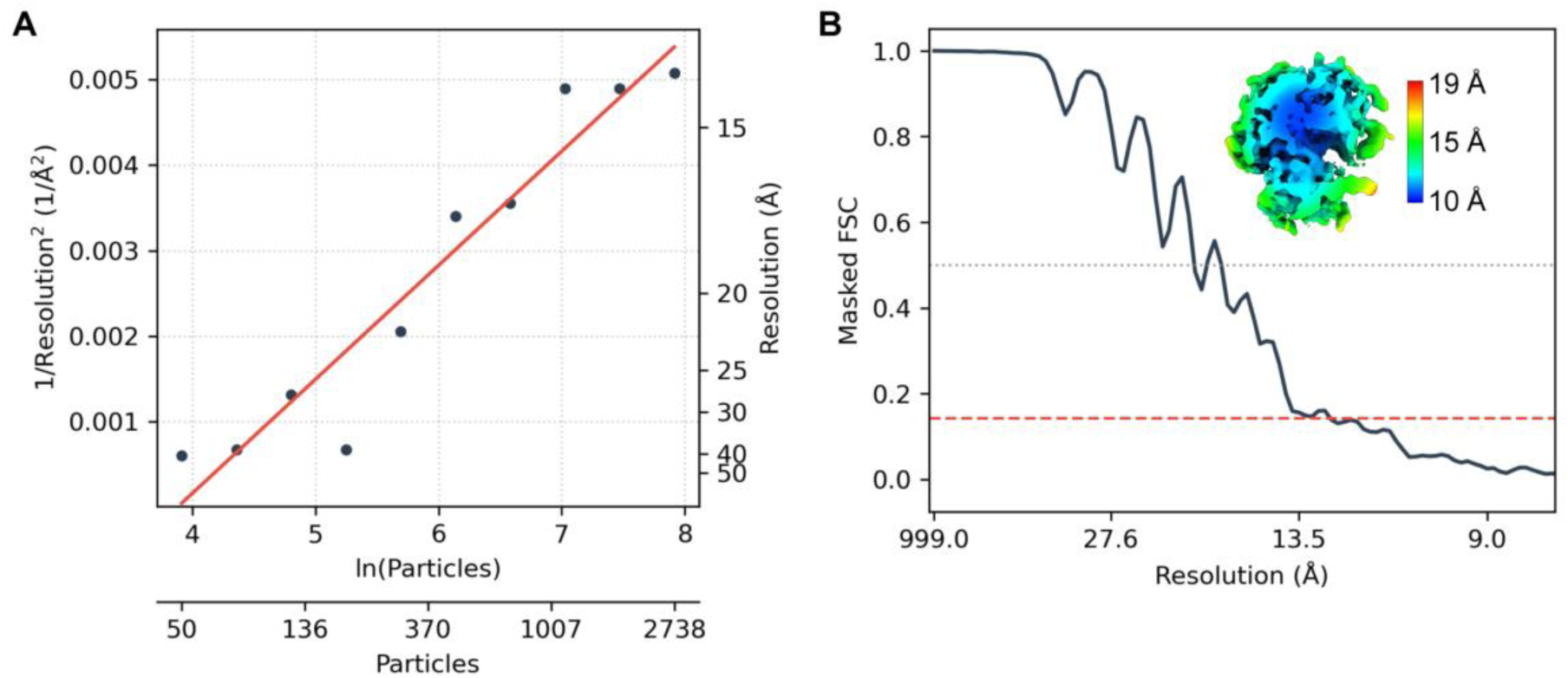
Resolution assessment of subtomogram averaging. a) Rosenthal–Henderson plot showing the gold-standard masked FSC resolution (as inverse squared) as a function of the natural logarithm of the number of particles. The fitted Rosenthal-Henderson B-factor is 1,502 Å^2^. b) Gold-standard masked FSC curve from the final RELION refinement. The dashed red line marks the 0.143 FSC criterion used to estimate the global resolution. The inset shows the final subtomogram average colored by local resolution as estimated by RELION post-processing, with the color scale indicating resolution values in Å.

3.3. *Subtomogram averaging results from sublimated lamellae*

Further evidence supporting the persistence of vitreous ice after sublimation is demonstrated by the preservation of the high resolution signal resulting from subtomogram averaging. We performed subtomogram averaging of the yeast 80s ribosome and estimated the Rosenthal-Henderson B-factor, resulting in a value of 1502 Å^2^ [25] (Fig. 5a). The gold-standard FSC curve (Fig. 5b) indicated a global resolution of 13.5 Å. The achieved resolution is comparable to previously published results by Tuijtel et al. [34] for thick lamellae obtained using a similar number of particles (182-255 nm lamella thickness; B-factor: −158Å^2^; ∼3000 particles; resolution: ∼9 Å for M-refinements Bin1: 1.2 Å/pix). Considering the low number of particles (∼2700), lamella thickness (287 ± 33 nm), and the high defocus (−6 µm), this resolution demonstrates the feasibility of high-resolution subtomogram averaging after sublimation.

3.4. *Fast-sublimation supplemental experiment*

While effective, sublimation conditions were found to be highly sensitive to thermal control. A supplementary experiment using a more rapid temperature ramp at a higher peak temperature led to visible pitting and surface damage, resulting in compromised lamella quality (Fig. S1). Although these lamellae also showed no signs of devitrification, they had extensive curtaining, rips, and even holes resulting from milling the damaged sample surface. It is highly possible that the warmer temperature and fast ramp rate from this experiment, similar to −95°C temperatures used in deep-etching experiments [17], led to etching away the ice between the small yeast cells.

Subsequently when lamellae were milled, these pores resulted in holes in the lamellae. Because the fast ramp condition also reached warmer temperatures (−97°C vs. −104°C), the individual contributions of ramp rate versus peak temperature cannot be isolated from the present data. These findings underscore the importance of careful sublimation parameter tuning. A slower, more gradual sublimation protocol using the minimum temperature required for sublimation was essential to maintain surface integrity. Table 1 denotes recommended sublimation settings used in this study, as well as experimental notes for each parameter.

**Table 1.** Troubleshooting and Parameters.

| **Parameter** | **Recommended Settings** | **Notes/Pitfalls** |
| --- | --- | --- |
| Temperature ramp rate | 7°C h<sup>-1</sup> (-118°C → -104°C) | Faster → pitting and rips |
| Sublimation temperature | -113°C to -104°C | Warmer experiment (-97°C) had pitting and rips |
| Observation interval | SEM every 5 min; FIB every 10-15 min | Stop when no ice visible |
| Cooling gas flow | 145 mg s<sup>-1</sup> → 200 mg s<sup>-1</sup> | Ensures rapid re-vitrification |

3.5. *Alternative surface cleaning methods*

Taken together, we show that sublimation can be implemented using standard cryo-FIB-SEM instrumentation without additional hardware, offering a practical and cost-effective approach to improving lamellae preparation. Aside from sublimation, one of the only other techniques used to remove surface contamination from cryo samples involves manually brushing the specimen with a cryo-compatible paintbrush under liquid nitrogen [9]. While this can be effective in some cases, it is mechanically invasive, operator-dependent, and difficult to standardize. Other surface cleaning strategies, such as ion polishing or plasma treatment [35,36], are only applicable to room temperature FIB milling and are not suitable for vitrified biological specimens. In contrast, sublimation provides a cryo-compatible, visually-guided step that enhances reproducibility and reduces milling artefacts. When appropriately controlled, it can be safely integrated into cryo-ET workflows and may serve as a useful addition to existing sample preparation protocols.

## 4. Conclusion

Our findings establish in-situ sublimation as a practical and effective method for improving cryo-lamellae preparation in cryo-ET workflows for HPF yeast waffle cells, without compromising ice quality. Broader applicability across plunge-frozen cells and HPF tissue samples will be evaluated in future work. By removing surface ice prior to milling, this approach enhances lamella uniformity and reduces curtaining artefacts, while preserving the vitrified state of the sample. These findings challenge longstanding assumptions about sublimation and demonstrate that, when properly controlled, it can be safely and reproducibly integrated into standard cryo-EM workflows. This method may be particularly valuable for high-pressure frozen samples, methods such as correlative light and electron microscopy (CLEM) involving multiple transfers between instruments, and other workflows where surface contamination is common. Future work should evaluate whether sublimation can be used on polished lamellae to remove surface ice accumulated during storage or fluorescence imaging (e.g., super-resolution) prior to cryo-ET imaging.

## Data Availability

The cryo-EM density maps for the 80S ribosome have been deposited in the Electron Microscopy Data Bank (EMDB) under accession codes EMD-56752.

## Acknowledgments

All data was collected and processed at the Karolinska Institutet 3D-EM facility.

## Author Contributions: CRediT

**Amy Bondy:** Conceptualization, Investigation, Writing, and Visualization. **Genis Valentin-Gese:** Formal analysis, Writing-Review & Editing, Visualization. **Thomas Thersleff:** Formal analysis, Writing-Review & Editing, Visualization. **B. Martin Hällberg:** Conceptualization, Writing-Review & Editing, Visualization, Supervision.

## Declaration of Generative AI Use

During the preparation of this work the authors used Large Language Models in order to improve language and readability. After using this tool, the authors reviewed and edited the content and take full responsibility for the content of the published article.

## Highlights

- In-chamber sublimation improves cryo-FIB sample preparation
- Controlled in-chamber sublimation reduces surface ice and FIB curtaining artefacts
- Milled lamellae show no devitrification during cryo-electron tomography

