## Supplemental_Information for "In-Chamber Sublimation: A Practical Approach for Mitigating Ice and Curtaining in Cryo-Electron Tomography Lamellae Preparation"

| **Table S1. Surface roughness (RMS squared) of SEM images* from Figure 2 during slow-sublimation experiment.** | |
|---|---|
| **Sublimation Time** | **Roughness (RMS)** ± standard error |
| $T_s$ = 0 | 0.065 ± 0.002 |
| $T_s$ = 15 | 0.060 ± 0.001 |
| $T_s$ = 30 | 0.056 ± 0.002 |
| $T_s$ = 45 | 0.048 ± 0.001 |

*All SEM images were normalized and filtered before roughness was calculated.

| **Table S2. M refinement parameters** | | | | | |
|---|---|---|---|---|---|
| **Geometry** | | | **Tilt series** | | **CTF** |
| M refinement | Image warp grid | Particle poses | Stage angles | Volume warp grid | Defocus |
| 1 | 3x3 | ✓ | ✓ | 3x3x2x8 | |
| 2 | 6x6 | ✓ | ✓ | 6x6x8x8 | |
| 3 | 6x6 | ✓ | ✓ | 6x6x8x8 | ✓ |

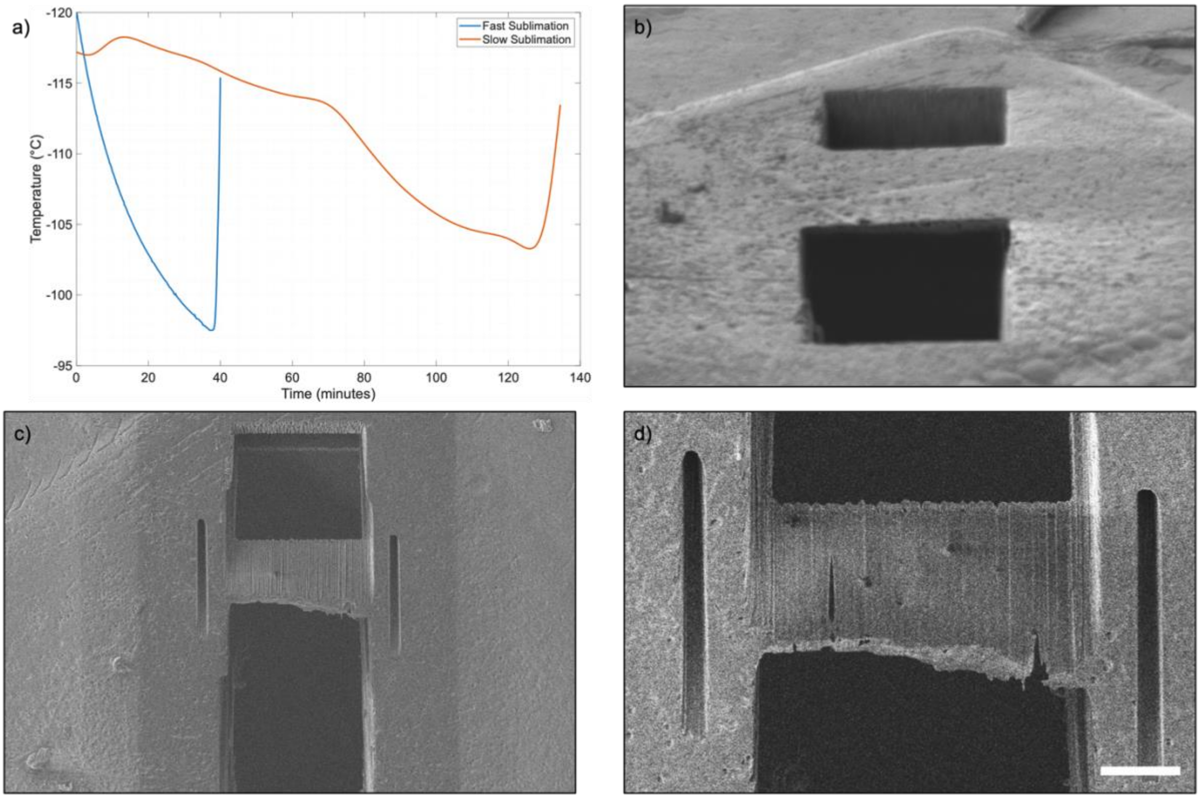


Figure S1. a) Experimental temperature curves showing sublimation of surface ice on samples during two different experiments. The red line denotes a slower sublimation rate and colder temperatures (corresponding with experimental data depicted in Figures 2- 5 in the main text), while the blue curve shows an experiment with a faster sublimation rate that reached warmer temperatures and resulted in a damaged surface. Faster sublimation rates appeared to cause “pitting” of the surface leading to holes and surface roughness that resulted in damaged lamellae visible after b) trench milling; c) rough milling); and d) thinning.
